# Plant G x Microbial E: Plant genotype interaction with soil bacterial community shapes rhizosphere composition during invasion

**DOI:** 10.1101/2024.04.19.590333

**Authors:** Mae Berlow, Miles Mesa, Mikayla Creek, Jesse Duarte, Elizabeth Carpenter, Brandon Phinizy, Krikor Andonian, Katrina M Dlugosch

## Abstract

It is increasingly recognized that different genetic variants can uniquely shape their microbiomes. Invasive species often evolve in their introduced ranges, but little is known about the potential for microbial associations to evolve during invasion as a result. We investigated invader genotype (G) and microbial environment (E) interactions in *C. solstitialis* (yellow starthistle), a Eurasian plant that is known to have evolved novel genotypes, and to have altered microbial interactions, in its severe invasion of California, USA. We conducted an experiment in which native and invading genotypes were inoculated with native and invaded range soil microbial communities. We used amplicon sequencing to characterize rhizosphere bacteria in both the experiment and the field soils from which they were derived. We found that bacterial diversity is higher in invaded soils, but that invading genotypes accumulated a lower diversity of bacteria and unique microbial composition in experimental inoculations, relative to native genotypes. Associations with potentially beneficial Streptomycetaceae were particularly interesting, as these were more abundant in the invaded range and accumulated on invading genotypes. Thus variation in microbial associations of invaders was driven by the interaction of G and E, and microbial communities appear to change in composition along with host evolution during invasion.

## INTRODUCTION

Introduced species that proliferate in their new environments are able to do so as a result of a suite of environmental and biological interactions (Sakai et al. 2001; Pearson et al. 2018). Biotic interactions are thought to be a particularly important means by which an introduced species could become invasive, because of opportunities for escape from enemies (i.e. the Enemy Release Hypothesis; (Keane and Crawley 2002; Engelkes et al. 2008; Colautti et al. 2004; Franks et al. 2008; Mitchell et al. 2010) or gain of mutualisms (Reinhart and Callaway 2006; Toby Kiers et al. 2010; van Kleunen, Bossdorf, and Dawson 2018; Sheng et al. 2022) in an introduced environment. Altered interactions with enemies or mutualists could have immediate benefits to invaders, but also might allow introduced populations to evolve traits that increase invasiveness, in response to increased resource availability from mutualists and/or relaxed selection for defense from enemies (Mesa and Dlugosch 2020; Blumenthal et al. 2009). While the potential for invaders to adapt in response to differences in species interactions in this way has received considerable attention (e.g. the pervasive Evolution of Increased Competitive Ability Hypothesis; (Blossey and Notzold 1995; Callaway et al. 2022), it is also the case that invader evolution (for any reason, adaptive or otherwise) might itself alter species interactions. Whether altered species interactions arise from a novel environment (‘E’) or from invader evolution (i.e. genotype, ‘G’), or their interaction (‘GxE’) is largely unexplored, though the contributions of these effects would have important consequences for understanding the mechanisms of invasions and predicting further spread (e.g. see (Huang, Lankau, and Peng 2018; Lankau et al. 2009).

For invasive plants, a critical source of both enemy and mutualist interactions is the soil microbial community (Dawkins et al. 2022; Rout and Callaway 2012; Callaway et al. 2004; Torchin and Mitchell 2004; Colautti et al. 2004; Agrawal et al. 2005; Kulmatiski et al. 2008; van der Putten et al. 2013; Faillace, Lorusso, and Duffy 2017; Dawson and Schrama 2016). Some invasive plants perform better with soil microbial communities from invaded ranges, suggesting that immediate interactions with soil microbes could facilitate invasion (Reinhart et al. 2003; Callaway et al. 2004; Mitchell et al. 2006; Engelkes et al. 2008; Maron et al. 2014; Andonian and Hierro 2011) but see (Shelby et al. 2016). Invaders could also benefit from differences in plant-soil feedbacks that accumulate over time (Levine et al. 2006). Plants experience negative feedbacks when a host-specific pathogen accumulates in the soil and suppresses their growth (Bever 2003, 2002), or positive feedbacks if the same dynamic occurs with a mutualist (Uddin et al. 2021), and invading species have been observed to benefit from enhanced positive or reduced negative feedbacks in soils from their introduced ranges (Uddin et al. 2021; Maron et al. 2014; McGinn et al. 2018; Meng et al. 2024; Suding et al. 2013; van der Putten et al. 2013). Invaders might also benefit from interactions with a simplified microbial interaction network in a novel environment, allowing invaders to be more efficient in exploiting mutualists or defending against pathogens (Colautti and Lau 2015; Mesa and Dlugosch 2020).

Regardless of whether microbial pathogens or mutualists differ between native and invaded ranges, we know very little about whether invaders that undergo evolutionary change over the course of their invasion interact with soil microbes differently than native genotypes. It is well documented in the fields of plant disease and agriculture that the genotype of a plant can be tightly correlated with the presence or relative abundance of specific microbes in and around their tissues (Yue et al. 2024; Schweitzer et al. 2008; Aira et al. 2010). Plant genotypes can control microbial communities through differences in their resulting phenotype, via traits like root and leaf morphology (King et al. 2021), exudate production (Saunders and Kohn 2009; Bailey et al. 2005), and interaction between plant phenotype and environment (e.g. root morphology) (Lareen, Burton, and Schäfer 2016). Plant genotypes can determine the amount or type of compounds they produce in defense against microbes (Van der Ent et al. 2008), or as incidental secondary metabolites that nonetheless impact microbial communities associating with their tissues (Voges et al. 2019). All of these genetically-based phenotypic differences have been shown to control plant association with individual pathogens or mutualists, as well as the plant-associated microbial community makeup (Saunders and Kohn 2009; Bailey et al. 2005; Ahlholm et al. 2002). The strong existing evidence that plant genotype affects microbial associations suggests that changes in invader genotype could lead to changes in microbial associations, which could then play a role in invader success and proliferation.

Indeed, invaders are well known to experience rapid evolution of novel genotypes for a variety of reasons. Introduced populations can adapt to their new environments and the process of colonization (Williams, Snyder, and Levine 2016; Williams, Kendall, and Levine 2016; Szűcs et al. 2017), but they may also experience non-adaptive genetic changes through several mechanisms, including initial and serial founder events, multiple introductions and admixture of previously isolated subpopulations, hybridization with other species, stimulation of transposable element activity, and/or the revealing of cryptic genetic variation in a novel environment (Katrina M. Dlugosch, Anderson, et al. 2015; K. M. Dlugosch and Parker 2008; Lee 2002; Bock et al. 2015; Colautti and Lau 2015; Prentis et al. 2008). Such broad opportunities for genetic change will create opportunities for altered species interactions, particularly with microbial communities that are sensitive to variation in plant growth and chemistry.

We test whether invader evolution alters microbial interactions in the invasive annual forb yellow starthistle (*Centaurea solstitialis*, L.; Asteraceae). This species is introduced and highly invasive in grasslands of North and South America (Gerlach 1997). Genotypes in invasive populations are known to have evolved to grow larger and produce more seeds than native genotypes (Widmer et al. 2007; Eriksen et al. 2012; Katrina M. Dlugosch, Alice Cang, et al. 2015). Invading plants also appear to benefit from more favorable (reduced deleterious) interactions with the soil microbial community in the invaded range (Andonian and Hierro 2011; Andonian et al. 2012, 2011), raising the possibility that the larger and more fecund invading genotypes have evolved to take advantage of release from microbial enemies or gain of mutualists (Lu-Irving et al. 2019). Field experiments with fungicide indicate that fungi are not the microbes responsible for more favorable soil interactions in the invasion, suggesting that bacterial communities might play an important role (Hierro et al. 2016). A previous survey of bacteria associated with *C. solstitialis* in the field found that bacterial diversity, including of known pathogenic groups, was lower on invading plants (Lu-Irving et al. 2019). These results suggest that invader genotypes might benefit from altered interactions with the microbial community, but it is not clear to what extent observed microbial differences are a cause or effect of the evolution of invader genotypes.

Here, we identify the roles of both microbial community environment, plant genotype, and their interaction in shaping plant-microbial associations in this system. We sample field soil from native and invaded ranges to identify the bacterial community available for plant interactions. We conduct a factorial greenhouse experiment, in which both native and invading *C. solstitialis* genotypes are grown in native and invaded range soil microbial communities, to identify which members of the soil community recruit to roots of each genotype in each range. We identify bacteria using 16S amplicon sequencing, and test for associations with genotype and soil source. We find that the interaction of both plant genotype and microbial environment influence specific microbial associations, and we find that previously-observed patterns of lower diversity of bacterial communities on invading plants are not inherent features of the environment as formerly hypothesized, but are instead the product of novel plant genotypes in the invasion.

## METHODS

### Study system

*Centaurea solstitialis* was introduced accidentally as a contaminant of alfalfa into South America in the 1600s and then North America in the 1800s (Gerlach 1997; DiTomaso and Healy 2007). It is an obligately outcrossing diploid plant that has four distinct native genetic subpopulations across Eurasia (M. Sun 1997; Eriksen et al. 2014; Barker et al. 2017; Irimia et al. 2017). Invading populations in California are genetically derived from the native subpopulation in western Europe (Barker *et al*. 2017). In both California and western Europe, *C. solstitialis* is an annual plant that grows as a rosette with a taproot through winter and spring, then bolts and flowers throughout the summer (Hierro et al. 2009; Katrina M. Dlugosch, Alice Cang, et al. 2015; Widmer et al. 2007).

### Soil and seed sampling

Soils used in this study were collected during dry summer conditions in August 2018 from four invaded sites in California, USA and four native sites in southern Spain and France (Supporting Information Table S1). At each site, a 30 m linear sampling transect was established through a *C. solstitialis* patch, and a second 30 m sampling transect was established outside of the patch, parallel to the first and separated by ∼5 m. A soil sample was collected every two meters for a total of 15 samples per transect. At each sampling point, a 18 mm diameter soil core was used to sample the topmost 5-10 cm of soil from three adjacent cores (<10 cm apart), which were combined into a single sample in a sealed plastic bag. Larger bulk soil collections (to be used for experimental inoculum) were made near each transect, both inside and outside of *C. solstitialis* patches. For these bulk collections, 1 L plastic bags were filled with soil collected from the top 5-10 cm of soil. Gloves and tools were sterilized between soil samples by wiping with 70% isopropanol. Seeds were collected from the plant closest to each meter along the transect through the *C. solstitialis* plants, for a total of 30 collections per site.

Soil collections were cleaned and homogenized for DNA sampling and use as inoculum. Large rocks and organic particles were removed from each sample manually with sterilized forceps in a biosafety cabinet, and the resulting sample sifted through a sterilized 2 mm sieve. Soil aggregates that did not pass through the sieve were ground gently with a mortar and pestle to break up the aggregates until all soil particles could pass through the 2mm sieve. The sample was stirred to homogenize, and 250 mg was weighed and collected in a 2 ml microfuge tube for DNA extraction.

### Experimental inoculations

Native and invaded range plant genotypes were grown in soils inoculated with native and invaded range live soils from the bulk collections made outside of *C. solstitialis* patches at same sites (Supporting Information Table S1) in a fully factorial greenhouse experiment. All seeds were germinated on the surface of sterile soil in petri dishes, in a greenhouse set to a maximum of 21℃ day and 10℃ night at the University of Arizona (Tucson, AZ, USA) College of Agriculture and Life Sciences greenhouses in February 2019. One week after germinating, seedlings were potted in individual Deepots**™** (410mL; D25L from Stuewe & Sons, Oregon USA) in sterile soil (50:50 mix of sand and high porosity soil; HP Pro-Mix**™,** Quebec, CA) and inoculated with a surface application of 15 mL of live soil slurry from soils collected in the same locations, or a sterile soil control. Slurries were prepared by adding 150 mL of sterile water to 10 mL of soil, mixing, and filtering through sterile cotton gauze.

Plants were maintained in the greenhouse until harvest at approximately two months of age. Sterile water was provided through daily misting by a reverse osmosis irrigation system (Evolution-RO, Hydro-Logic Inc., Port Washington, NY, USA). After one month, plants were fertilized every two weeks using autoclaved Hoagland’s solution (⅛ strength Hoagland Complete Medium, BioWorld, Dublin, OH, USA). At harvest, plants were removed from their pots and the upper 2 to 5 cm of the taproot was collected, together with accompanying lateral roots (excess soil was brushed or shaken off). Root samples were placed in individual 50 ml tubes containing 25 ml of sterile wash solution (45.9 mM NaH2PO4, 61.6 mM Na2HPO4, 0.1% Tween 20). Tubes were shaken by hand for 1 minute. Root samples were then removed and stored on ice in tubes containing 10 ml of wash solution until further processing, stored on ice during transport and then refrigerated at 4°C. Ectorhizosphere washes were centrifuged at 2,200 g at 4°C for 15 min, supernatants were discarded, and pellets were air-dried and stored at 20°C until DNA extraction.

### DNA extraction and amplicon sequencing

Microbial DNA was extracted from soil samples using the DNeasy PowerSoil Pro kit (Qiagen, Hilden, Germany). A blank sample of nanopure water was also included in extractions to record any contamination. Resulting double-stranded DNA concentration was quantified using a Qubit fluorometer (Broad Range kit, Invitrogen, Waltham MA, USA).

The 16s rRNA region was amplified using a 2-step polymerase chain reaction (PCR) as in Lu-Irving et. al. (2019). Target-specific PCR was conducted by creating a 25.1 μL reaction mixture using 1 μL of microbial DNA, 1.3 μL of 515-F primer and 1.3 μL of 806-R primer, 12.5 μL of Phusion Flash master mix (Thermo Scientific, Waltham MA, USA), and 9 μL PCR grade water. Reaction mixtures were placed in an Eppendorf Mastercycler thermal cycler starting with 98℃ for 10 seconds, then 25 cycles of this sequence: 98℃ for 1 second, 78℃ for 5 seconds, 57℃ for 5 seconds, 72℃ for 15 seconds, 72℃ for 1 minute (Lundberg et al. 2013). Step 1 PCR products were visualized on an agarose gel to determine whether PCR was successful before a second PCR step was used to incorporate sequencing adapters onto the ends of the amplified PCR products. Step 2 reaction mixtures were created using 1uL of PCR product, 12.5 μL Phusion Flash master mix, and 0.75 μL of a unique barcoded primer combination provided by the University of Idaho’s iBEST Genomic Resources Core. Our PCR program ran at 98℃ for 10 seconds, then 10 cycles of: 98℃ for 1 second, 78℃ for 5 seconds, 51℃ for 5 seconds, 72℃ for 15 seconds, 72℃ for 1 minute. Barcoded amplicons were quantified by Qubit fluorometry, pooled in equimolar amounts, cleaned using a MinElute kit (Qiagen, Hilden, Germany), and submitted to the University of Idaho’s iBEST Genomic Resources Core for quality control and sequencing with Illumina MiSeq 350 bp pair-end sequencing.

### Analyses

Microbial metabarcoding data was processed and analyzed in Qiime 2 version 2019.10 (Bolyen et al. 2019), and additional analyses were carried out in R (R Core Team 2019), as detailed below. Scripts for processing sequences and replicating all analyses are available on GitHub (https://anonymous.4open.science/r/Plant-G-x-Microbial-E-F8D3/README.md). Sequences were denoised in Qiime 2 using the Divisive Amplicon Denoising Algorithm (DADA2) to remove sequence errors and trim primers (Rosen et al. 2012). Next, sequences were aligned, and a phylogeny was generated using FastTree, rooted at the midpoint (Price, Dehal, and Arkin 2010). Sequences were grouped at the level of amplicon sequence variants (ASVs, 100% similarity) and taxonomy was assigned using the Silva database, version 138 (Quast et al. 2013). ASVs assigned to non-bacterial kingdoms were filtered out for the purposes of our analyses.

Bacterial alpha diversity was measured as richness calculated by the R package ‘vegetarian’ (Jost 2007) and Shannon’s Diversity Index calculated by Qiime2 after rarefying to 1000 sequences per sample (Bolyen et al. 2019). To test for the effects of soil range (native vs. invaded) on alpha diversity metrics, we used two-sided t-tests in the stats package in R (R Core Team 2019). To test for effects of experimental treatments on alpha diversity metrics, we used linear models that included effects of seed genotype (native vs. invader), soil range (native vs. invading), germination date of the plant (i.e. its age at harvest), experimental block in the greenhouse, and the interaction between seed genotype and soil range. Non-significant interaction terms (P > 0.1) were not included in the final models. Linear models and Tukey’s post hoc tests were conducted using the stats package in R (R Core Team 2019). To determine which bacterial taxa were differentially abundant between sample types and treatment groups we used Linear Discriminant Analysis Effect Size (LEfSe, (Segata et al. 2011).

Bacterial community beta diversity was measured as unweighted and weighted UniFrac distances, a dissimilarity measure that accounts for phylogenetic relatedness (Lozupone et al. 2011). Unweighted UniFrac distance accounts for information about presence/absence of ASVs and can be thought of as community membership, while weighted UniFrac distance also accounts for relative abundances and can be thought of as community structure. We used PERMANOVA conducted in the R package vegan (Oksanen 2015) to test whether beta diversity distances were predicted by fixed effects of range (native vs. invaded), and sites nested within range (including samples both inside and outside of YST patches).

To test whether the bulk soil collections (used as inocula for the greenhouse experiment) were representative of the nearby soil bacterial diversity, we compared the distances between each bulk soil sample and the soil cores from the same site with the distances between that same bulk soil sample and all the soil cores from other sites using paired t-tests in R.

## RESULTS

We sequenced a total of 351 samples, yielding 16,654,396 paired reads (mean = 44,059, SD = 35,066, see Table S2 for sequence and ASV counts for each sample). Sequences are available on the NCBI Sequence Repository (SUB13812121).

### Field environment: Soils

Across our native and invaded range field soil samples we identified 29 phyla, including 294 families of bacteria. Eleven phyla made up 99% of sequences present (Figure 1, Table S3). The most abundant phyla across all samples were Actinobacteriota (an average of 33% of invaded soil bacterial communities, 41% of native), Proteobacteria (22% of invaded, 20% of native), and Acidobacteriota (14% of invaded, 18% of native).

**Figure 1.**
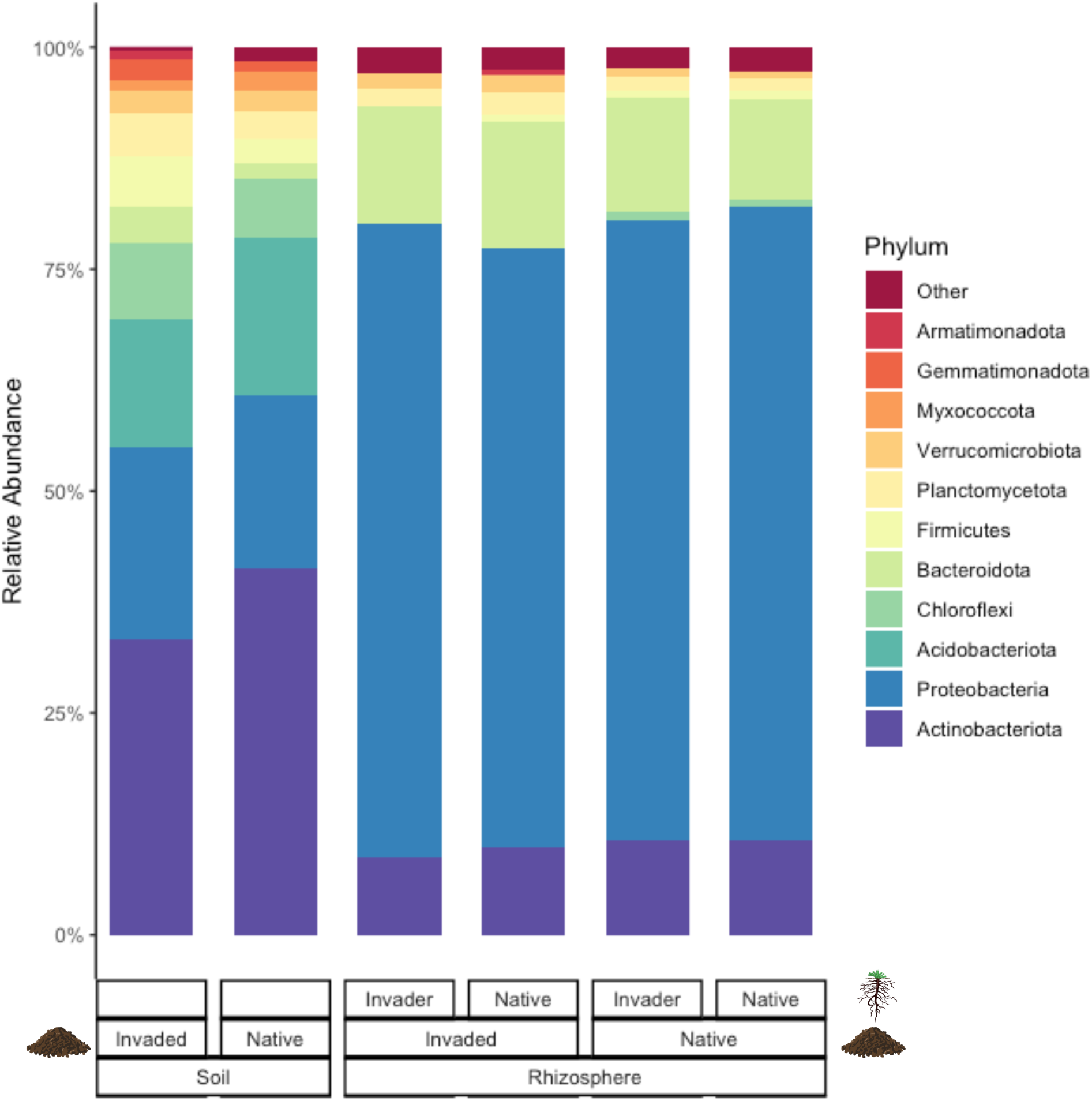
Relative abundance of bacterial phyla in field soil samples, grouped by range (invaded or native) and rhizosphere samples (root wash) grouped by plant genotype (invader or native) within range. All phyla shown constitute at least 1% of at least one sample for soil, and at least 0.6% of at least one sample for rhizosphere. Relative abundance of each phylum averaged across samples for each group.

Bacterial taxa differed between the native and invaded range soils at both the phylum and family level. Ten phyla were differentially abundant: Actinobacteriota, Acidobacteriota, and Myxococcota were more abundant in native range soils, and Proteobacteria, Armatimonadota, Gemmatimonadota, Planctomycetota, Chloroflexi, Bacteriodota, and Firmicutes were more abundant in invaded range soils (LDA effect size >2, LEfSe; Figure S1). Nineteen bacterial families were more abundant in native range soils, and 55 families were more abundant in invaded range soils (LDA effect size >2, LEfSe; Table S4).

The distribution of soil bacterial diversity also differed between the ranges. Invaded soils had higher alpha diversity than native soils by both metrics (richness *P* = 0.035, Shannon *P* =0.002, *t*-test; Figure 2a-b). In terms of beta diversity, both range (native or invaded) and site had a significant effect on community membership and community structure (all *P* < 0.0001, PERMANOVA; Table 1; Figure 3).

**Figure 2.**
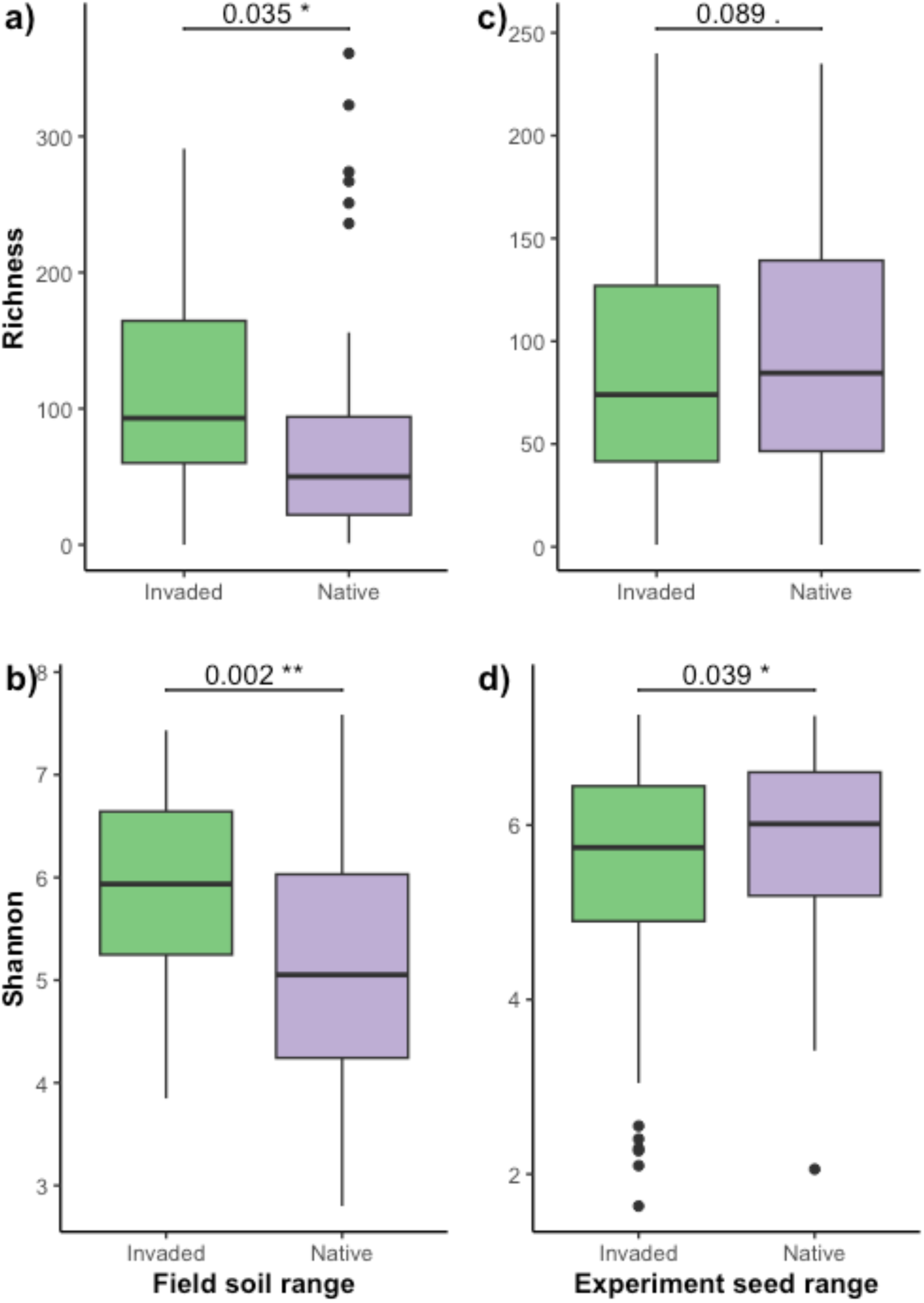
Box plots show two measures of alpha diversity (a & c richness, b & d Shannon’s diversity index) for field soil samples (a & b) in invaded range soils (California, USA) and native range soils (Spain and France) and for native and invader yellow starthistle (*C. solstitialis*) genotypes grown (c & d).

**Figure 3.**
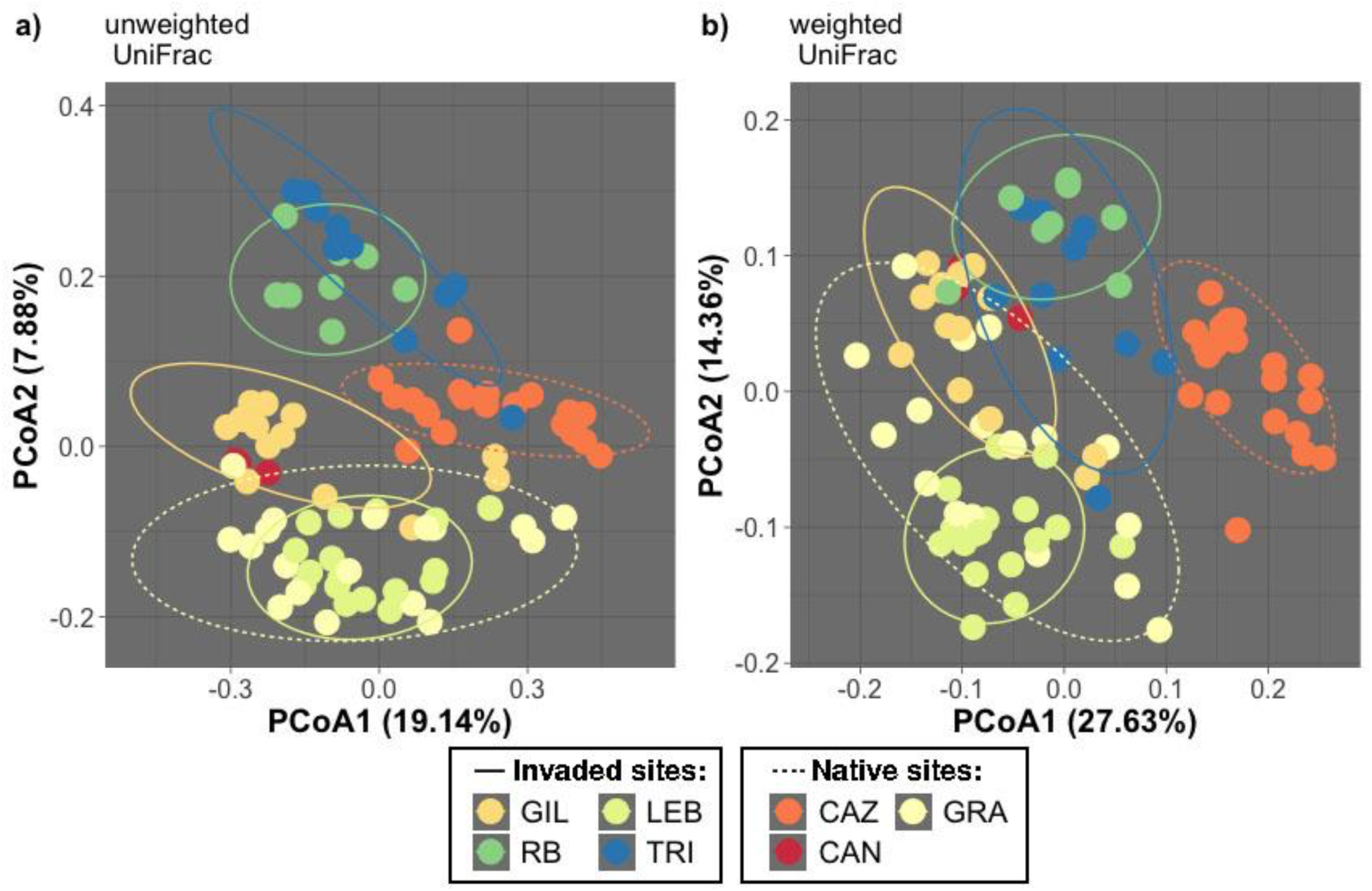
Principle coordinates ordination of **a)** weighted UniFrac distances (community structure) and **b)** unweighted uniFrac distances (community membership) of field soil samples. Variation explained by each axis is shown in parentheses. Each data point represents one soil sample. Ellipses represent 90% confidence intervals.

**Table 1.** PERMANOVA results for two measures of beta diversity (weighted and unweighted UniFrac).

| predictor |  | df | Pseudo-F | P |
| --- | --- | --- | --- | --- |
| *unweighted UniFrac:* |  |  |  |  |
| Range |  | 1 | 5.48 | 0.0001 |
| Site with/in range |  | 6 | 5.49 | 0.0001 |
| *weighted UniFrac:* |  |  |  |  |
| Range |  | 1 | 10.15 | 0.0001 |
| Site with/in range |  | 6 | 12.51 | 0.0001 |

We found that the distances between the bulk soils used as inocula and the soil cores from the same site were significantly smaller than the distances between the bulk soils and soil cores from external sites (unweighted unifrac *P* = 0.005, weighted unifrac *P* = 0.0001, paired *t*-test, Figure S2). In other words, bulk soils used as inocula in the greenhouse experiment were more similar to the soil of the site they were from than to any of our other sites, and were thus representative of those sites for use in and interpretation of our greenhouse experiments.

### Experimental inoculation: Rhizosphere

Across our rhizosphere samples we identified 23 phyla and 352 families of bacteria. Eight phyla made up 99% of sequences present (Figure 1, Table S5). The most abundant phyla were Proteobacteria (an average of 70% across samples), Bacteroidota (13%), and Actinobacteriota (10%). There were no phyla that differed between native and invading *C. solstitialis* genotypes within either native or invaded soil treatments, or between native and invader plant genotypes overall (all LDA effect sizes <2, LEfSe).

At the bacterial family level, the interaction of the source of soil microbial communities and plant genotype shaped rhizosphere communities. For invaded range soil inocula, seven bacterial families were differentially abundant, including 6 families that were more abundant on native range plants and one that was more abundant on invaders (LDA effect size > 2, LEfSe; Figure 4). For native range soil inocula, five families differed, with two families more abundant on native range plants and three that were more abundant on invaders (LDA effect size > 2, LEfSe; Figure 4). Only the family Micropepsaceae was differentially abundant between plant genotypes in both native and invaded soil treatments, in which it was more abundant on native genotypes in both treatments.

**Figure 4.**
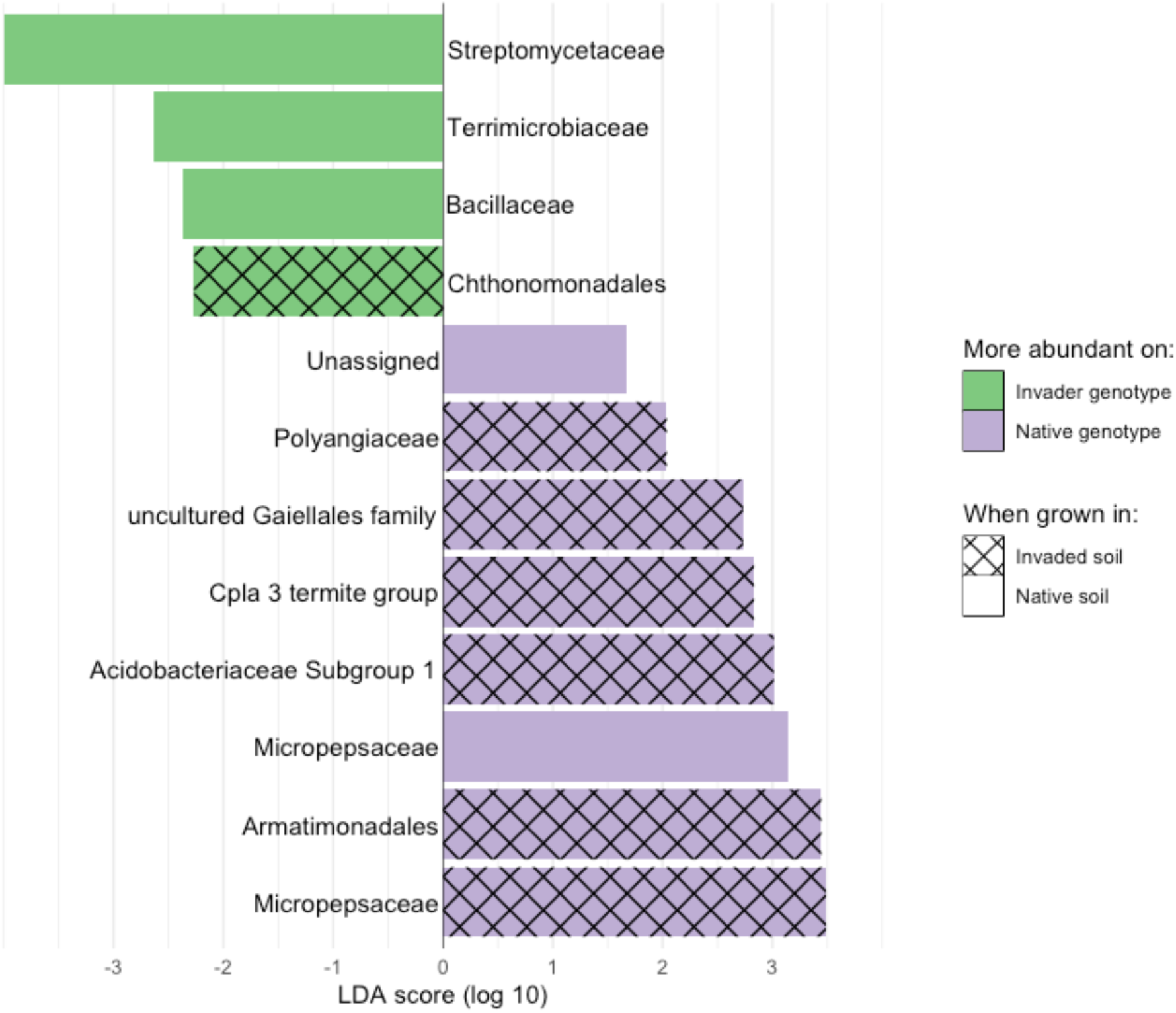
Bacterial families that are differentially abundant in rhizosphere bacterial communities between invader and native *C. solstitialis* genotypes when grown with invaded and native microbial communities. Results from linear discriminant analysis effect size (LEfSe). Note: Micropepsaceae was the only family found to be differentially abundant between genotypes in both invaded and native soil.

Contrary to patterns in field soils, alpha diversity (richness and Shannon’s H) was higher for bacterial communities on native *C. solstitialis* genotypes than for communities on invader plant genotypes (Figure 2c-d, Table S6). There was not a significant difference in alpha diversity between communities derived from native vs. invaded range soils, nor an interaction between soil range and plant genotype (Table S6).

## DISCUSSION

Our goal in this study was to test for the effects of plant evolution and its interaction with the soil microbial environment on plant-microbial associations during invasion. We found that both genotype and environment shaped microbial associations. We also found that previously-observed patterns of plant-associated microbial diversity in this invasion (Lu-Irving et al. 2019) were not a product of microbial communities available in the environment. Instead, genotype drove the pattern of microbial diversity, such that invading plant genotypes accumulated a lower diversity of rhizosphere bacteria than native genotypes, despite invaded range soils being overall higher in microbial diversity.

Amplicon sequencing of field soils indicated that bacterial community environments differed significantly between ranges, and among sites within ranges. Differences between native and invaded microbial environments were evident at both the phylum and family levels. For both taxonomic levels most differentially abundant taxa were higher in the invaded range soils than the native range soils (e.g. 55 families were more abundant in the invaded range, versus 19 that were elevated in the native range). Consistent with these differences, alpha diversity (both richness and Shannon diversity) was higher in the invaded range soils. Soil microbial diversity and composition are known to vary geographically for a variety of reasons, including soil physical characteristics (particularly pH), climatic factors, land use and disturbance, and plant species composition (Fierer and Jackson 2006; Nottingham et al. 2018; Chu et al. 2020). Our collection sites were typically roadsides adjacent to agricultural fields, with cover of *C. solstitialis* and European grasses, and climatically similar environments (Barker et al. 2017), in both ranges. Nevertheless, significant site effects within each range indicate that microbial communities were sensitive to subtleties of site characteristics, and the distribution of some microbial taxa (see below) suggest that native range sites might experience more drought stress.

Using these same soils as inocula in our experiment, we found that invading genotypes associated with a lower diversity of bacteria than did native genotypes, regardless of the source of the soil inoculum. This indicates that plant genotypes in this system shape the microbial communities on their roots, and that invaders are not only experiencing novel interactions, their evolution is shaping these interactions. In their observational study of *C. solstitialis*-associated microbial communities in the field, Lu-Irving and colleagues (2019) noted that rhizosphere microbial diversity was positively correlated with the genetic diversity of plants at a site within each range, consistent with an important effect of genotype in the field. This relationship could not explain differences in microbial diversity between ranges, however, because plant genetic diversity *per se* does not differ substantially between the native and invaded range in this system (Lu-Irving et al. 2019; Barker et al. 2017). Instead, our experiment indicates that it is divergence between native and invading plant genotypes (rather than differences in their population genetic diversity) that explains lower rhizosphere diversity on plants in the invasion.

We found no effect of soil community environment on alpha diversity of root-associated microbes in our experiment, but we did find effects of both soil environment and plant genotype on the abundance of specific taxa in the rhizosphere. Below we discuss known functions of several of these taxa, but we emphasize that their relationships to *C. solstitialis* are unknown in all cases and must be investigated further. While it is difficult to generalize the functions of many bacterial taxa (especially at higher taxonomic ranks), and some are entirely unknown, current understanding of these groups suggests that additional research on their functional roles in this system would be informative.

In possibly our most important finding regarding differentially abundant taxa, we found that Streptomycetaceae were more abundant in invaded range field soils, and were more abundant on invader genotypes when grown in the native range soils (in which this group was less abundant, suggesting preferential accumulation). This association between Streptomycetaceae and the invaded range, and the apparent accumulation of these bacteria on invading genotypes, is potentially important because the Streptomycetaceae are widely associated with the promotion of plant growth (Viaene et al. 2016). Their benefits appear to be both direct effects on growth (as ‘biofertilizers’) and indirect effects through their suppression of plant pathogens (the Streptomycetaceae are also responsible for the majority of human antibiotics) (Olanrewaju and Babalola 2019). If members of this family have a similar positive relationship with invading *C. solstitialis*, this could explain several observed advantages to invaders in this system, including increased invader growth (Katrina M. Dlugosch, Alice Cang, et al. 2015), reduced negative plant-soil interactions in the invaded range (Andonian and Hierro 2011), and potentially the decrease in diversity of microbes on invader roots, given the antimicrobial properties of Streptomycetaceae (Lu-Irving et al. 2019).

Native genotypes also had a set of interactions that mirrored the invader relationships with Streptomycetaceae. These were taxa that were more abundant in native soils, and also more abundant on native genotypes when grown in the invaded soils in which these same groups were more depauperate (again suggesting preferential accumulation). These groups have a variety of potential functions of interest, and include an uncultured Gaiellales family (a group that has been associated with plant colonizers, and may regulate organic and fatty acids in soils (Ye et al. 2021; L. Sun et al. 2022)); a family in Subgroup 1 of the Acidobacteriaceae (a diverse group of oligotrophs abundant in soils, some of which have adapted to specific dry soil conditions; (Kielak et al. 2016; Sikorski et al. 2022)), a family in the CPla-3 termite group (a more highly specialized group in the soil, often in acidic environments; (Fickling et al. 2024)), and the family Polyangiaceae (a group of predatory social bacteria, among the most common predators of bacteria in the soil, with the potential to structure the diversity of the entire microbial community (Petters et al. 2021; Z. Wu et al. 2022; Thakur and Geisen 2019)).

The most notable association for native plant genotypes, however, appeared to be with the family Micropepsaceae, which was more abundant in invaded range soils, but was overrepresented on native genotypes grown in either invaded or native range soils. This was the only family to be overrepresented on a genotype across both soil environments. The Micropepsaceae has been found to be an indicator taxon associated with specific plant species or site conditions in other studies (D. Wu et al. 2023; Gschwend et al. 2022). For studies that compared different environmental manipulations, the family was associated with early phases of restoration (grasslands; (Barber et al. 2023)), early responses to agricultural planting (tobacco; (Cao et al. 2022)), and response to cold treatments of plants (lettuce; (Persyn et al. 2022)), all conditions in which a system had been perturbed recently. In one study, the presence of Micropepsaceae was also associated with reduced suppression of pathogens (Yu et al. 2022). If native genotypes are exposed to more pathogens when associating with this family (perhaps during resource acquisition after disturbance), this might favor diversion of resources from growth info defenses, which has been hypothesized to explain the smaller size of native plant genotypes within *C. solstitialis* (Lu-Irving et al. 2019).

The association patterns of two families suggest that the native environment might be more stressful for plants, particularly in terms of drought. While native and invaded range climatic niches are generally similar in these areas (Barker et al. 2017), the families Bacillaceae and Terrimicrobiaceae were more abundant in native range soils. Bacillaceae is characterized by the ability to form endospores that are resistant to environmental extremes, including drought (Mandic-Mulec, Stefanic, and van Elsas 2016). The Terrimicrobiaceae have been observed to be enriched for G3P (Glycerol-3-phosphate) transport-related genes, which are associated with plant-microbe interactions that promote drought tolerance of plants (Styer et al. 2024). Both families were also more abundant on invader genotypes in native soils, which is notable because previous research has found that invader genotypes are less drought tolerant, and have suggested that invaders have adapted to an environment with lower competition for limiting resources, primarily summer water (Katrina M. Dlugosch, Alice Cang, et al. 2015).

Finally, two groups that are more common in invaded range soils could be of interest for their differential associations with each genotype growing in that soil inoculum. Taxa in the Armatimonadales (a group of putatively generalist consumers of organic compounds; (Kato et al. 2022; Fickling et al. 2024) have also been associated with invasions of *Amaranthus palmeri* in China (Zhang et al. 2023), but were more abundant on native genotypes of *C. solstitialis*. In contrast, taxa in the Chthonomonadales (a group associated with decomposition of glycine substrates; (Bhatnagar, Peay, and Treseder 2018) were more abundant on invader genotypes.

Further work is needed to identify how *C. solstitialis* interactions with these specific bacteria, as well as other components of their microbial communities (i.e. fungi and viruses), affect host plant fitness across ranges. Microbial interactions could be responsible for natural selection that has driven evolution of invaders in this system, given that root microbiomes can have large effects on host plant performance (Bai et al. 2022; Trivedi et al. 2020). A study of 15 annual plant species in California grasslands (not including *C. solstitialis*), found that soil microbial communities generated large fitness differences among species, suggesting that microbially-driven selection could be strong (Kandlikar गौरव कांडलकर et al. 2021).

Invaders will also evolve to adapt to other aspects of their new environments, and are expected to evolve increases in traits associated with invasiveness itself (Williams, Snyder, and Levine 2016; Williams, Kendall, and Levine 2016; Weiss-Lehman, Hufbauer, and Melbourne 2017; Szűcs et al. 2017; Perkins et al. 2013; Katrina M. Dlugosch, Alice Cang, et al. 2015; Gioria et al. 2023; McGaughran et al. 2024). Such trait changes could be synergistic with more favorable microbial interactions, for example the evolution of increased root investments for resource acquisition could also help plants take advantage of beneficial microbes (Dawson 2015). The influence of host evolution on microbial interactions during invasion does not appear to have been investigated previously, however. There is some evidence that plant-microbial interactions have changed over time during invasion, which could result from plant evolutionary change. For example, older invading populations of *Solidago canadensis* were found to have increased positive microbial interactions (and competitive ability) in a common garden experiment, suggesting that these changes were genetically-based (Oduor et al. 2022). In contrast, the evolution of aboveground herbivore interactions in plant invasions has attracted longstanding interest (Mesa and Dlugosch 2020), and recent reviews find abundant evidence for the evolution of these interactions, though not consistently to the advantage or disadvantage of the invader (Gruntman and Segev 2024; Yi et al. 2024). Our work demonstrates that belowground microbial interactions should also be expected to evolve during invasion, as a product of both the resident microbial community, and the genetic composition of the host. How these interactions alter fitness and invasiveness over time, and how they interact with available genetic variation in introduced populations, will be important avenues of further research.

## Reference

Agrawal, Anurag A., Peter M. Kotanen, Charles E. Mitchell, Alison G. Power, William Godsoe, and John Klironomos. 2005. “Enemy Release? An Experiment with Congeneric Plant Pairs and Diverse above- and Belowground Enemies.” Ecology 86 (11): 2979–89.

Ahlholm, Jouni U., Marjo Helander, Janne Henriksson, Mary Metzler, and Kari Saikkonen. 2002. “Environmental Conditions and Host Genotype Direct Genetic Diversity of Venturia Ditricha, a Fungal Endophyte of Birch Trees.” Evolution; International Journal of Organic Evolution 56 (8): 1566–73.

Aira, Manuel, María Gómez-Brandón, Cristina Lazcano, Erland Bååth, and Jorge Domínguez. 2010. “Plant Genotype Strongly Modifies the Structure and Growth of Maize Rhizosphere Microbial Communities.” Soil Biology & Biochemistry 42 (12): 2276–81.

Andonian, Krikor, and José L. Hierro. 2011. “Species Interactions Contribute to the Success of a Global Plant Invader.” Biological Invasions 13 (12): 2957–65.

Andonian, Krikor, José L. Hierro, Liana Khetsuriani, Pablo I. Becerra, Grigor Janoyan, Diego Villareal, Lohengrin A. Cavieres, Laurel R. Fox, and Ragan M. Callaway. 2012. “Geographic Mosaics of Plant-Soil Microbe Interactions in a Global Plant Invasion.” Journal of Biogeography 39 (3): 600–608.

Andonian, Krikor, José L. Hierro, Liana Khetsuriani, Pablo Becerra, Grigor Janoyan, Diego Villarreal, Lohengrin Cavieres, Laurel R. Fox, and Ragan M. Callaway. 2011. “Range-Expanding Populations of a Globally Introduced Weed Experience Negative Plant-Soil Feedbacks.” PLoS One 6 (5): e20117.

Bai, Bo, Weidong Liu, Xingyu Qiu, Jie Zhang, Jingying Zhang, and Yang Bai. 2022. “The Root Microbiome: Community Assembly and Its Contributions to Plant Fitness.” Journal of Integrative Plant Biology 64 (2): 230–43.

Bailey, Joseph K., Ron Deckert, Jennifer A. Schweitzer, Brian J. Rehill, Richard L. Lindroth, Catherine Gehring, and Thomas G. Whitham. 2005. “Host Plant Genetics Affect Hidden Ecological Players: Links among Populus, Condensed Tannins, and Fungal Endophyte Infection.” Canadian Journal of Botany. Journal Canadien de Botanique 83 (4): 356–61.

Barber, Nicholas A., Desirae M. Klimek, Jennifer K. Bell, and Wesley D. Swingley. 2023. “Restoration Age and Reintroduced Bison May Shape Soil Bacterial Communities in Restored Tallgrass Prairies.” FEMS Microbiology Ecology 99 (3). 10.1093/femsec/fiad007.

Barker, B. S., K. Andonian, S. M. Swope, D. G. Luster, and K. M. Dlugosch. 2017. “Population Genomic Analyses Reveal a History of Range Expansion and Trait Evolution across the Native and Invaded Range of Yellow Starthistle (Centaurea Solstitialis).” Molecular Ecology 26 (4): 1131–47.

Bever, James D. 2002. “Negative Feedback within a Mutualism: Host-Specific Growth of Mycorrhizal Fungi Reduces Plant Benefit.” Proceedings. Biological Sciences / The Royal Society 269 (1509): 2595–2601.

Bever, James D. 2003. “Soil Community Feedback and the Coexistence of Competitors: Conceptual Frameworks and Empirical Tests.” The New Phytologist 157 (3): 465–73.

Bhatnagar, Jennifer M., Kabir G. Peay, and Kathleen K. Treseder. 2018. “Litter Chemistry Influences Decomposition through Activity of Specific Microbial Functional Guilds.” Ecological Monographs 88 (3): 429–44.

Blossey, Bernd, and Rolf Notzold. 1995. “Evolution of Increased Competitive Ability in Invasive Nonindigenous Plants: A Hypothesis.” The Journal of Ecology 83 (5): 887–89.

Blumenthal, Dana, Charles E. Mitchell, Petr Pysek, and Vojtech Jarosík. 2009. “Synergy between Pathogen Release and Resource Availability in Plant Invasion.” Proceedings of the National Academy of Sciences of the United States of America 106 (19): 7899–7904.

Bock, D. G., C. Caseys, R. G. Cousens, M. A. Hahn, S. M. Heredia, S. Hubner, K. G. Turner, K. D. Whitney, and Loren H. Rieseberg. 2015. “What We Still Don’t Know about Invasion Genetics.” Molecular Ecology 24: 2277–97.

Bolyen, Evan, Jai Ram Rideout, Matthew R. Dillon, Nicholas A. Bokulich, John Chase, Emily K. Cope, Kestrel Gorlick, et al. 2019. “Reproducible, Interactive, Scalable and Extensible Microbiome Data Science Using QIIME 2 (Nature Biotechnology, (2019), 37, 8, (852-857), 10.1038/s41587-019-0209-9).” Nature Biotechnology 37 (8): 852–57.

Callaway, Ragan M., Jacob E. Lucero, José L. Hierro, and C. J. Lortie. 2022. “The EICA Is Dead? Long Live the EICA!” Ecology Letters 25 (10): 2289–2302.

Callaway, Ragan M., Giles C. Thelen, Alex Rodriguez, and William E. Holben. 2004. “Soil Biota and Exotic Plant Invasion.” Nature 427 (6976): 731–33.

Cao, Yi, Zhi-Xiao Yang, Dong-Mei Yang, Ning Lu, Shi-Zhou Yu, Jian-Yu Meng, and Xing-Jiang Chen. 2022. “Tobacco Root Microbial Community Composition Significantly Associated With Root-Knot Nematode Infections: Dynamic Changes in Microbiota and Growth Stage.” Frontiers in Microbiology 13 (February): 807057.

Chu, Haiyan, Gui-Feng Gao, Yuying Ma, Kunkun Fan, and Manuel Delgado-Baquerizo. 2020. “Soil Microbial Biogeography in a Changing World: Recent Advances and Future Perspectives.” mSystems 5 (2). 10.1128/mSystems.00803-19.

Colautti, Robert I., and Jennifer A. Lau. 2015. “Contemporary Evolution during Invasion: Evidence for Differentiation, Natural Selection, and Local Adaptation.” Molecular Ecology 24 (9): 1999–2017.

Colautti, Robert I., Anthony Ricciardi, Igor A. Grigorovich, and Hugh J. MacIsaac. 2004. “Is Invasion Success Explained by the Enemy Release Hypothesis?” Ecology Letters 7 (8): 721– 33.

Dawkins, K., J. Mendonca, O. Sutherland, and N. Esiobu. 2022. “A Systematic Review of Terrestrial Plant Invasion Mechanisms Mediated by Microbes and Restoration Implications.” American Journal of Plant Sciences 13 (02): 205–22.

Dawson, Wayne. 2015. “Release from Belowground Enemies and Shifts in Root Traits as Interrelated Drivers of Alien Plant Invasion Success: A Hypothesis.” Ecology and Evolution 5 (20): 4505–16.

Dawson, Wayne, and Maarten Schrama. 2016. “Identifying the Role of Soil Microbes in Plant Invasions.” The Journal of Ecology 104 (5): 1211–18.

DiTomaso, Joseph M., and E. A. Healy. 2007. Weeds of California and Other Western States. Oakland, CA: University of California Department of Agriculture and Natural Resources.

Dlugosch, Katrina M., F. Alice Cang, Brittany S. Barker, Krikor Andonian, Sarah M. Swope, and Loren H. Rieseberg. 2015. “Evolution of Invasiveness through Increased Resource Use in a Vacant Niche.” Nature Plants 1 (6): 15066.

Dlugosch, Katrina M., Samantha R. Anderson, Joseph Braasch, F. Alice Cang, and Heather D. Gillette. 2015. “The Devil Is in the Details: Genetic Variation in Introduced Populations and Its Contributions to Invasion.” Molecular Ecology 24: 2095–2111.

Dlugosch, K. M., and I. M. Parker. 2008. “Founding Events in Species Invasions: Genetic Variation, Adaptive Evolution, and the Role of Multiple Introductions.” Molecular Ecology 17 (1): 431–49.

Engelkes, Tim, Elly Morriën, Koen J. F. Verhoeven, T. Martijn Bezemer, Arjen Biere, Jeffrey A. Harvey, Lauren M. McIntyre, Wil L. M. Tamis, and Wim H. van der Putten. 2008. “Successful Range-Expanding Plants Experience Less above-Ground and below-Ground Enemy Impact.” Nature 456 (7224): 946–48.

Eriksen, Renée L., Theodora Desronvil, José L. Hierro, and Rick Kesseli. 2012. “Morphological Differentiation in a Common Garden Experiment among Native and Non-Native Specimens of the Invasive Weed Yellow Starthistle (Centaurea Solstitialis).” Biological Invasions 14 (7): 1459–67.

Eriksen, Renée L., José L. Hierro, Özkan Eren, Krikor Andonian, Katalin Török, Pablo I. Becerra, Daniel Montesinos, Liana Khetsuriani, Alecu Diaconu, and Rick Kesseli. 2014. “Dispersal Pathways and Genetic Differentiation among Worldwide Populations of the Invasive Weed Centaurea Solstitialis L. (Asteraceae).” PloS One 9 (12): e114786.

Faillace, Cara A., Nicholas S. Lorusso, and Siobain Duffy. 2017. “Overlooking the Smallest Matter: Viruses Impact Biological Invasions.” Ecology Letters EarlyView (February). 10.1111/ele.12742.

Fickling, Nicole W., Catherine A. Abbott, Joel E. Brame, Christian Cando-Dumancela, Craig Liddicoat, Jake M. Robinson, and Martin F. Breed. 2024. “Light-Dark Cycles May Influence in Situ Soil Bacterial Networks and Diurnally-Sensitive Taxa.” Ecology and Evolution 14 (2): e11018.

Fierer, Noah, and Robert B. Jackson. 2006. “The Diversity and Biogeography of Soil Bacterial Communities.” Proceedings of the National Academy of Sciences 103 (3): 626–31.

Franks, Steven J., Paul D. Pratt, F. Allen Dray, and Ellen L. Simms. 2008. “Selection on Herbivory Resistance and Growth Rate in an Invasive Plant.” The American Naturalist 171 (5): 678– 91.

Gerlach, John D. 1997. “How the West Was Lost: Reconstructing the Invasion Dynamics of Yellow Starthistle and Other Plant Invaders of Western Rangelands and Natural Areas.” California Exotic Pest Plant Council Symposium Proceedings 3: 67–72.

Gioria, Margherita, Philip E. Hulme, David M. Richardson, and Petr Pyšek. 2023. “Why Are Invasive Plants Successful?” Annual Review of Plant Biology 74 (May): 635–70.

Gruntman, Michal, and Udi Segev. 2024. “Effect of Residence Time on Trait Evolution in Invasive Plants: Review and Meta-Analysis.” Working Paper Series 91 (February): 99–124.

Gschwend, Florian, Martin Hartmann, Johanna Mayerhofer, Anna-Sofia Hug, Jürg Enkerli, Andreas Gubler, Reto G. Meuli, Beat Frey, and Franco Widmer. 2022. “Site and Land-Use Associations of Soil Bacteria and Fungi Define Core and Indicative Taxa.” FEMS Microbiology Ecology 97 (12). 10.1093/femsec/fiab165.

Hierro, José L., Özkan Eren, Liana Khetsuriani, Alecu Diaconu, Katalin Török, Daniel Montesinos, Krikor Andonian, et al. 2009. “Germination Responses of an Invasive Species in Native and Non-Native Ranges.” Oikos 118 (4): 529–38.

Hierro, José L., Liana Khetsuriani, Krikor Andonian, Özkan Eren, Diego Villarreal, Grigor Janoian, Kurt O. Reinhart, and Ragan M. Callaway. 2016. “The Importance of Factors Controlling Species Abundance and Distribution Varies in Native and Non-Native Ranges.” Ecography EarlyView. 10.1111/ecog.02633.

Huang, Fangfang, Richard Lankau, and Shaolin Peng. 2018. “Coexistence via Coevolution Driven by Reduced Allelochemical Effects and Increased Tolerance to Competition between Invasive and Native Plants.” The New Phytologist 218 (1): 357–69.

Irimia, Ramona-Elena, Daniel Montesinos, Özkan Eren, Christopher J. Lortie, Kristine French, Lohengrin A. Cavieres, Gastón J. Sotes, José L. Hierro, Andreia Jorge, and João Loureiro. 2017. “Extensive Analysis of Native and Non-Native Centaurea Solstitialis L. Populations across the World Shows No Traces of Polyploidization.” PeerJ 5 (August): e3531.

Jost, Lou. 2007. “Partitioning Diversity into Independent Alpha and Beta Components.” Ecology 88 (10): 2427–39.

Kandlikar गौरव कां(डलकर, Gaurav S., Xinyi Yan 严心怡, Jonathan M. Levine, and Nathan J. B. Kraft. 2021. “Soil Microbes Generate Stronger Fitness Differences than Stabilization among California Annual Plants.” The American Naturalist 197 (1): E30–39.

Kato, Shingo, Sachiko Masuda, Arisa Shibata, Ken Shirasu, and Moriya Ohkuma. 2022. “Insights into Ecological Roles of Uncultivated Bacteria in Katase Hot Spring Sediment from Long-Read Metagenomics.” Frontiers in Microbiology 13 (November): 1045931.

Keane, Ryan M., and Michael J. Crawley. 2002. “Exotic Plant Invasions and the Enemy Release Hypothesis.” Trends in Ecology & Evolution 17 (4): 164–70.

Kielak, Anna M., Cristine C. Barreto, George A. Kowalchuk, Johannes A. van Veen, and Eiko E. Kuramae. 2016. “The Ecology of Acidobacteria: Moving beyond Genes and Genomes.” Frontiers in Microbiology 7 (May): 744.

King, William L., Caylon F. Yates, Jing Guo, Suzanne M. Fleishman, Ryan V. Trexler, Michela Centinari, Terrence H. Bell, and David M. Eissenstat. 2021. “The Hierarchy of Root Branching Order Determines Bacterial Composition, Microbial Carrying Capacity and Microbial Filtering.” Communications Biology 4 (1): 483.

Kleunen, Mark van, Oliver Bossdorf, and Wayne Dawson. 2018. “The Ecology and Evolution of Alien Plants.” Annual Review of Ecology, Evolution, and Systematics 49 (1): 25–47.

Kulmatiski, Andrew, Karen H. Beard, John R. Stevens, and Stephanie M. Cobbold. 2008. “Plant– soil Feedbacks: A Meta-Analytical Review.” Ecology Letters 11 (9): 980–92.

Lankau, Richard A., Victoria Nuzzo, Greg Spyreas, and Adam S. Davis. 2009. “Evolutionary Limits Ameliorate the Negative Impact of an Invasive Plant.” Proceedings of the National Academy of Sciences 106: 15362–67.

Lareen, Andrew, Frances Burton, and Patrick Schäfer. 2016. “Plant Root-Microbe Communication in Shaping Root Microbiomes.” Plant Molecular Biology 90 (6): 575–87.

Lee, Carol Eunmi. 2002. “Evolutionary Genetics of Invasive Species.” Trends in Ecology & Evolution 17 (8): 386–91.

Levine, Jonathan M., Elizaveta Pachepsky, Bruce E. Kendall, Stephanie G. Yelenik, and Janneke Hille Ris Lambers. 2006. “Plant-Soil Feedbacks and Invasive Spread.” Ecology Letters 9 (9): 1005–14.

Lozupone, Catherine, Manuel E. Lladser, Dan Knights, Jesse Stombaugh, and Rob Knight. 2011. “UniFrac: An Effective Distance Metric for Microbial Community Comparison.” The ISME Journal 5 (2): 169–72.

Lu-Irving, Patricia, Julia G. Harenčár, Hailey Sounart, Shana R. Welles, Sarah M. Swope, David A. Baltrus, and Katrina M. Dlugosch. 2019. “Native and Invading Yellow Starthistle (Centaurea Solstitialis) Microbiomes Differ in Composition and Diversity of Bacteria.” mSphere 4 (2). 10.1128/mSphere.00088-19.

Lundberg, Derek S., Scott Yourstone, Piotr Mieczkowski, Corbin D. Jones, and Jeffery L. Dangl. 2013. “Practical Innovations for High-Throughput Amplicon Sequencing.” Nature Methods 10 (10): 999–1002.

Mandic-Mulec, Ines, Polonca Stefanic, and Jan Dirk van Elsas. 2016. “Ecology of *Bacillaceae*.” In The Bacterial Spore, 59–85. Washington, DC, USA: ASM Press.

Maron, John L., John Klironomos, Lauren Waller, and Ragan M. Callaway. 2014. “Invasive Plants Escape from Suppressive Soil Biota at Regional Scales.” The Journal of Ecology 102 (1): 19– 27.

McGaughran, Angela, Manpreet K. Dhami, Elahe Parvizi, Amy L. Vaughan, Dianne M. Gleeson, Kathryn A. Hodgins, Lee A. Rollins, et al. 2024. “Genomic Tools in Biological Invasions: Current State and Future Frontiers.” Genome Biology and Evolution 16 (1). 10.1093/gbe/evad230.

McGinn, Kevin J., Wim H. van der Putten, Philip E. Hulme, Natasha Shelby, Carolin Weser, and Richard P. Duncan. 2018. “The Influence of Residence Time and Geographic Extent on the Strength of Plant-Soil Feedbacks for Naturalised Trifolium.” Edited by Matthew Turnbull. The Journal of Ecology 106 (1): 207–17.

Meng, Yunhao, Xinze Geng, Ping Zhu, Xinfu Bai, Ping Zhang, Guangyan Ni, and Yuping Hou. 2024. “Enhanced Mutualism: A Promotional Effect Driven by Bacteria during the Early Invasion of Phytolacca Americana.” Ecological Applications: A Publication of the Ecological Society of America 34 (1): e2742.

Mesa, J. Miles, and Katrina M. Dlugosch. 2020. “The Evolution of Invasiveness: A Mechanistic View of Trade-offs Involving Defenses.” American Journal of Botany 107 (7): 953–56.

Mitchell, Charles E., Anurag A. Agrawal, James D. Bever, Gregory S. Gilbert, Ruth A. Hufbauer, John N. Klironomos, John L. Maron, et al. 2006. “Biotic Interactions and Plant Invasions.” Ecology Letters 9 (6): 726–40.

Mitchell, Charles E., Dana Blumenthal, Vojtěch Jarošík, Emily E. Puckett, and Petr Pyšek. 2010. “Controls on Pathogen Species Richness in Plants’ Introduced and Native Ranges: Roles of Residence Time, Range Size and Host Traits.” Ecology Letters 13 (12): 1525–35.

Nottingham, Andrew T., Noah Fierer, Benjamin L. Turner, Jeanette Whitaker, Nick J. Ostle, Niall P. McNamara, Richard D. Bardgett, et al. 2018. “Microbes Follow Humboldt: Temperature Drives Plant and Soil Microbial Diversity Patterns from the Amazon to the Andes.” Ecology 99 (11): 2455–66.

Oduor, Ayub M. O., Michael Opoku Adomako, Yongge Yuan, and Jun-Min Li. 2022. “Older Populations of the Invader Solidago Canadensis Exhibit Stronger Positive Plant-Soil Feedbacks and Competitive Ability in China.” American Journal of Botany 109 (8): 1230–41.

Oksanen, Jari. 2015. “Multivariate Analysis of Ecological Communities in R: Vegan Tutorial.” R Documentation, 43.

Olanrewaju, Oluwaseyi Samuel, and Olubukola Oluranti Babalola. 2019. “Streptomyces: Implications and Interactions in Plant Growth Promotion.” Applied Microbiology and Biotechnology 103 (3): 1179–88.

Pearson, Dean E., Yvette K. Ortega, Özkan Eren, and José L. Hierro. 2018. “Community Assembly Theory as a Framework for Biological Invasions.” Trends in Ecology & Evolution 33 (5): 313– 25.

Perkins, T. Alex, Benjamin L. Phillips, Marissa L. Baskett, and Alan Hastings. 2013. “Evolution of Dispersal and Life History Interact to Drive Accelerating Spread of an Invasive Species.” Ecology Letters 16 (8): 1079–87.

Persyn, Antoine, Sonia Garcia Mendez, Stien Beirinckx, Sam De Meyer, Anne Willems, Caroline De Tender, and Sofie Goormachtig. 2022. “Digging into the Lettuce Cold-Specific Root Microbiome in Search of Chilling Stress Tolerance-Conferring Plant Growth-Promoting Bacteria.” Phytobiomes Journal, December. 10.1094/PBIOMES-07-22-0044-MF.

Petters, Sebastian, Verena Groß, Andrea Söllinger, Michelle Pichler, Anne Reinhard, Mia Maria Bengtsson, and Tim Urich. 2021. “The Soil Microbial Food Web Revisited: Predatory Myxobacteria as Keystone Taxa?” The ISME Journal 15 (9): 2665–75.

Prentis, Peter J., John R. U. Wilson, Eleanor E. Dormontt, David M. Richardson, and Andrew J. Lowe. 2008. “Adaptive Evolution in Invasive Species.” Trends in Plant Science 13 (6): 288– 94.

Price, Morgan N., Paramvir S. Dehal, and Adam P. Arkin. 2010. “FastTree 2--Approximately Maximum-Likelihood Trees for Large Alignments.” PloS One 5 (3): e9490.

Putten, Wim H. van der, Richard D. Bardgett, James D. Bever, T. Martijn Bezemer, Brenda B. Casper, Tadashi Fukami, Paul Kardol, et al. 2013. “Plant–soil Feedbacks: The Past, the Present and Future Challenges.” The Journal of Ecology 101 (2): 265–76.

Quast, Christian, Elmar Pruesse, Pelin Yilmaz, Jan Gerken, Timmy Schweer, Pablo Yarza, Jörg Peplies, and Frank Oliver Glöckner. 2013. “The SILVA Ribosomal RNA Gene Database Project: Improved Data Processing and Web-Based Tools.” Nucleic Acids Research. 10.1093/nar/gks1219.

R Core Team. 2019. “R: A Language and Environment for Statistical Computing. Version 3.6. 0. Vienna, Austria.” R Foundation for Statistical Computing, Vienna, Austria. https://www.R-project.org/.

Reinhart, Kurt O., and Ragan M. Callaway. 2006. “Soil Biota and Invasive Plants.” The New Phytologist 170 (3): 445–57.

Reinhart, Kurt O., Alissa Packer, Wim H. Van der Putten, and Keith Clay. 2003. “Plant-Soil Biota Interactions and Spatial Distribution of Black Cherry in Its Native and Invasive Ranges.” Ecology Letters 6 (12): 1046–50.

Rosen, Michael J., Benjamin J. Callahan, Daniel S. Fisher, and Susan P. Holmes. 2012. “Denoising PCR-Amplified Metagenome Data.” BMC Bioinformatics 13 (1): 1–16.

Rout, Marnie E., and Ragan M. Callaway. 2012. “Interactions between Exotic Invasive Plants and Soil Microbes in the Rhizosphere Suggest That ‘Everything Is Not Everywhere.’” Annals of Botany 110 (2): 213–22.

Sakai, A. K., Fred W. Allendorf, J. S. Holt, D. M. Lodge, J. Molofsky, K. A. With, S. Baughman, et al. 2001. “The Population Biology of Invasive Species.” Annual Review of Ecology and Systematics 32: 305–32.

Saunders, Megan, and Linda Myra Kohn. 2009. “Evidence for Alteration of Fungal Endophyte Community Assembly by Host Defense Compounds.” The New Phytologist 182 (1): 229–38.

Schweitzer, Jennifer A., Joseph K. Bailey, Dylan G. Fischer, Carri J. LeRoy, Eric V. Lonsdorf, Thomas G. Whitham, and Stephen C. Hart. 2008. “Plant-Soil Microorganism Interactions: Heritable Relationship between Plant Genotype and Associated Soil Microorganisms.” Ecology 89 (3): 773–81.

Segata, Nicola, Jacques Izard, Levi Waldron, Dirk Gevers, Larisa Miropolsky, Wendy S. Garrett, and Curtis Huttenhower. 2011. “Metagenomic Biomarker Discovery and Explanation.” Genome Biology 12 (6): R60.

Shelby, Natasha, Richard P. Duncan, Wim H. Putten, Kevin J. McGinn, Carolin Weser, and Philip E. Hulme. 2016. “Plant Mutualisms with Rhizosphere Microbiota in Introduced versus Native Ranges.” The Journal of Ecology 104 (5): 1259–70.

Sheng, Min, Christoph Rosche, Mohammad Al-Gharaibeh, Lorinda S. Bullington, Ragan M. Callaway, Taylor Clark, Cory C. Cleveland, et al. 2022. “Acquisition and Evolution of Enhanced Mutualism-an Underappreciated Mechanism for Invasive Success?” The ISME Journal 16 (11): 2467–78.

Sikorski, Johannes, Vanessa Baumgartner, Klaus Birkhofer, Runa S. Boeddinghaus, Boyke Bunk, Markus Fischer, Bärbel U. Fösel, et al. 2022. “The Evolution of Ecological Diversity in Acidobacteria.” Frontiers in Microbiology 13 (February): 715637.

Styer, Alex, Dean Pettinga, Daniel Caddell, and Devin Coleman-Derr. 2024. “Improving Rice Drought Tolerance through Host-Mediated Microbiome Selection.” bioRxiv. 10.1101/2024.02.03.578672.

Suding, Katharine N., William Stanley Harpole, Tadashi Fukami, Andrew Kulmatiski, Andrew S. MacDougall, Claudia Stein, and Wim H. van der Putten. 2013. “Consequences of Plant–soil Feedbacks in Invasion.” The Journal of Ecology 101 (2): 298–308.

Sun, Litao, Yu Wang, Dexin Ma, Linlin Wang, Xiaomei Zhang, Yiqian Ding, Kai Fan, et al. 2022. “Differential Responses of the Rhizosphere Microbiome Structure and Soil Metabolites in Tea (Camellia Sinensis) upon Application of Cow Manure.” BMC Microbiology 22 (1): 55.

Sun, Mei. 1997. “Population Genetic Structure of Yellow Starthistle (Centaurea Solstitialis), a Colonizing Weed in the Western United States.” Canadian Journal of Botany. Journal Canadien de Botanique 75 (9): 1470–78.

Szűcs, M., M. L. Vahsen, B. A. Melbourne, C. Hoover, C. Weiss-Lehman, and R. A. Hufbauer. 2017. “Rapid Adaptive Evolution in Novel Environments Acts as an Architect of Population Range Expansion.” Proceedings of the National Academy of Sciences of the United States of America 114 (51): 13501–6.

Thakur, Madhav P., and Stefan Geisen. 2019. “Trophic Regulations of the Soil Microbiome.” Trends in Microbiology 27 (9): 771–80.

Toby Kiers, E., Todd M. Palmer, Anthony R. Ives, John F. Bruno, and Judith L. Bronstein. 2010. “Mutualisms in a Changing World: An Evolutionary Perspective.” Ecology Letters 13 (12): 1459–74.

Torchin, Mark E., and Charles E. Mitchell. 2004. “Parasites, Pathogens, and Invasions by Plants and Animals.” Frontiers in Ecology and the Environment 2 (4): 183–90.

Trivedi, Pankaj, Jan E. Leach, Susannah G. Tringe, Tongmin Sa, and Brajesh K. Singh. 2020. “Plant-Microbiome Interactions: From Community Assembly to Plant Health.” Nature Reviews. Microbiology 18 (11): 607–21.

Uddin, Md N., Takashi Asaeda, Animesh Sarkar, Viraj P. Ranawakage, and Randall W. Robinson. 2021. “Conspecific and Heterospecific Plant–soil Biota Interactions of Lonicera Japonica in Its Native and Introduced Range: Implications for Invasion Success.” Plant Ecology 222 (12): 1313–24.

Van der Ent, Sjoerd, Bas W. M. Verhagen, Ronald Van Doorn, Daniel Bakker, Maarten G. Verlaan, Michiel J. C. Pel, Ruth G. Joosten, et al. 2008. “MYB72 Is Required in Early Signaling Steps of Rhizobacteria-Induced Systemic Resistance in Arabidopsis.” Plant Physiology 146 (3): 1293–1304.

Viaene, Tom, Sarah Langendries, Stien Beirinckx, Martine Maes, and Sofie Goormachtig. 2016. “Streptomyces as a Plant’s Best Friend?” FEMS Microbiology Ecology 92 (8). 10.1093/femsec/fiw119.

Voges, Mathias J. E. E. E., Yang Bai, Paul Schulze-Lefert, and Elizabeth S. Sattely. 2019. “Plant-Derived Coumarins Shape the Composition of an Arabidopsis Synthetic Root Microbiome.” Proceedings of the National Academy of Sciences of the United States of America 116 (25): 12558–65.

Weiss-Lehman, Christopher, Ruth A. Hufbauer, and Brett A. Melbourne. 2017. “Rapid Trait Evolution Drives Increased Speed and Variance in Experimental Range Expansions.” Nature Communications 8 (January): 14303.

Widmer, Timothy L., Fatiha Guermache, Margarita Yu Dolgovskaia, and Sergey Ya Reznik. 2007. “Enhanced Growth and Seed Properties in Introduced vs. Native Populations of Yellow Starthistle (Centaurea Solstitialis).” Weed Science 55 (5): 465–73.

Williams, Jennifer L., Bruce E. Kendall, and Jonathan M. Levine. 2016. “Rapid Evolution Accelerates Plant Population Spread in Fragmented Experimental Landscapes.” Science 353 (6298): 482–85.

Williams, Jennifer L., Robin E. Snyder, and Jonathan M. Levine. 2016. “The Influence of Evolution on Population Spread through Patchy Landscapes.” The American Naturalist 188 (1): 15–26.

Wu, Di, Hui Bai, Caihong Zhao, Mu Peng, Qi Chi, Yaping Dai, Fei Gao, Qiang Zhang, Minmin Huang, and Ben Niu. 2023. “The Characteristics of Soil Microbial Co-Occurrence Networks across a High-Latitude Forested Wetland Ecotone in China.” Frontiers in Microbiology 14 (March): 1160683.

Wu, Zhaojun, Yang Li, Hao Chen, Jixiang Rao, and Qingye Sun. 2022. “Effects of Straw Mulching on Predatory Myxobacterial Communities in Different Soil Aggregates under Wheat-Corn Rotation.” Environmental Science and Pollution Research International 29 (19): 29062–74.

Ye, Fei, Xiaoxiao Wang, Yu Wang, Shengjun Wu, Jiapeng Wu, and Yiguo Hong. 2021. “Different Pioneer Plant Species Have Similar Rhizosphere Microbial Communities.” Plant and Soil 464 (1): 165–81.

Yi, Jiahui, Jinlong Wan, Katja Tielbörger, Zhibin Tao, Evan Siemann, and Wei Huang. 2024. “Specialist Reassociation and Residence Time Modulate the Evolution of Defense in Invasive Plants: A Meta-Analysis.” Ecology 105 (3): e4253.

Yue, Hong, Xuming Sun, Tingting Wang, Ali Zhang, Dejun Han, Gehong Wei, Weining Song, and Duntao Shu. 2024. “Host Genotype-Specific Rhizosphere Fungus Enhances Drought Resistance in Wheat.” Microbiome 12 (1): 44.

Yu, Li, Haiyun Zi, Hongguang Zhu, Yangwenke Liao, Xia Xu, and Xiaogang Li. 2022. “Rhizosphere Microbiome of Forest Trees Is Connected to Their Resistance to Soil-Borne Pathogens.” Plant and Soil 479 (1): 143–58.

Zhang, Mei, Cong Shi, Xueying Li, Kefan Wang, Zhenlu Qiu, and Fuchen Shi. 2023. “Changes in the Structure and Function of Rhizosphere Soil Microbial Communities Induced by Amaranthus Palmeri Invasion.” Frontiers in Microbiology 14 (March): 1114388.

